# SARS-CoV-2 ORF9b Antagonizes Type I and III Interferons by Targeting Multiple Components of RIG-I/MDA-5-MAVS, TLR3-TRIF, and cGAS-STING Signaling Pathways

**DOI:** 10.1101/2020.08.16.252973

**Authors:** Lulu Han, Meng-Wei Zhuang, Yi Zheng, Jing Zhang, Mei-Ling Nan, Pei-Hui Wang, Chengjiang Gao

**Affiliations:** Key Laboratory of Infection and Immunity of Shandong Province, Department of Immunology, School of Basic Medical Sciences, Cheeloo College of Medicine, Shandong University, Jinan 250012, China; Advanced Medical Research Institute, Cheeloo College of Medicine, Shandong University, Jinan 250012, China; Suzhou Research Institute, Shandong University, Shandong University, Suzhou, Jiangsu 215123, China

**Keywords:** SARS-CoV-2, COVID-19, ORF9b, antiviral immunity, IFNs

## Abstract

Severe acute respiratory syndrome corona-virus 2 (SARS-CoV-2), the etiologic agent of the coronavirus disease 2019 (COVID-19), has a catastrophic effect on human health and society. Clinical findings indicated that the suppression of innate antiviral immunity, especially the type I and III interferon (IFN) production, contributes to the pathogenesis of COVID-19. However, how SARS-CoV-2 evades antiviral immunity still needs further investigations. Here, we reported that the open reading frame 9b (ORF9b) protein encoded by the SARS-CoV-2 genome inhibits the activation of type I and III IFN response by targeting multiple molecules of innate antiviral signaling pathways. SARS-CoV-2 ORF9b impaired the induction of type I and III IFNs by Sendai virus or the dsRNA mimic poly (I:C). SARS-CoV-2 ORF9b inhibits the activation of type I and III IFNs induced by the components of cytosolic dsRNA-sensing pathways of RIG-I/MDA5-MAVS signaling, including RIG-I, MDA-5, MAVS, TBK1, and IKKε rather than IRF3-5D, the active form of IRF3. SARS-CoV-2 ORF9b also suppressed the induction of type I and III IFNs by TRIF and STING, the adaptor protein of endosome RNA-sensing pathway of TLR3-TRIF signaling and the adaptor protein of cytosolic DNA-sensing pathway of cGAS-STING signaling, respectively. Mechanistically, SARS-CoV-2 ORF9b protein interacts with RIG-I, MDA-5, MAVS, TRIF, STING, TBK1, and prevents TBK1 phosphorylation, thus impeding the phosphorylation and nuclear trans-localization of IRF3 activation. Overexpression of SARS-CoV-2 ORF9b facilitates the replication of the vesicular stomatitis virus. Therefore, SARS-CoV-2 ORF9b negatively regulates antiviral immunity, thus, facilitate virus replication. This study contributes to our understanding of the molecular mechanism of how SARS-CoV-2 impaired antiviral immunity and providing an essential clue to the pathogenesis of COVID-19.

## Introduction

Coronaviruses belong to a family of enveloped, positive-sense, single-stranded RNA viruses with 27.6–31 kb genome in size, which is associated with various natural hosts, such as amphibians, birds, and mammals.^1^ Severe acute respiratory syndrome coronavirus 2 (SARS-CoV-2), which causes coronavirus disease 2019 (COVID-19), is a novel emerging coronavirus that is spreading globally and may be lethal to humans and other animals, posing significant threats to public health worldwide.^2-4^ In the 21st century, SARS-CoV-1, Middle East respiratory syndrome human (MERS-CoV), and SARS-CoV-2 have emerged in succession in the human population and may cause severe pulmonary disease with acute respiratory distress syndrome and systemic inflammation and maybe fatality. Besides SARS-CoV-1, MERS-CoV, and SARS-CoV-2, other four coronaviruses that can infect human, including human coronavirus (HCoV)-229E, HCoV-NL63, HCoV-OC43, and HCoV-HKU1, which, in most condition, only cause the common cold via the infection of the human upper respiratory tract, but in children, elderly, and immunocompromised patients, they also lead to severe and even fatal diseases.^5^ The genome of SARS-CoV-2 is about 30 kb in length encoding 14 putative open reading frames, including the large replicase genes expressing two replicative polyproteins (pp1a and pp1ab) that would be cleaved into NSP1-16 by viral proteases, the structural genes expressing spike (S), membrane (M), envelop (E), and nucleocapsid (N) protein, and the accessory genes expressing ORF3a, ORF3b, ORF6, ORF7a, ORF7b, ORF8, ORF9b, ORF9c, and ORF10.^6^ Although the accessory proteins of coronaviruses are not essential for viral replication and virion assembly, they contribute to the virulence by affecting the virus releasement, stability, and pathogenesis.^7^ Up to date, the function of SARS-CoV-2 accessory proteins in immune evasion still need to be addressed.

Although the dysregulation of the immune system with suppressed antiviral immunity and elevated inflammatory responses caused by SARS-CoV-2 infection contribute to the pathogenesis of COVID-19,^8-13^ how SARS-CoV-2 was recognized by innate immunity is currently unknown since its recently emerging. dsRNA, produced by many viruses during replication, is a common viral pathogen-associated molecular patterns (PAMPs) assumed for immune sensing by pattern recognition receptors (PRRs).^14^ The cytosolic retinoic acid-inducible gene (RIG)-I-like receptors (RLRs) and the endosomal toll-like receptors (TLRs) recognize dsRNAs from intermediates generated during viral replication, resulting in the serial activation of signaling cascades to induce the production of type I and III IFNs.^14,15^ The coronaviruses have a homologous genome, the similar replication intermediates, and a same life cycle; thus, it seems that SARS-CoV-2 can be similarly recognized by the RNA sensors as other coronaviruses to elicit immune responses.^16^ MDA5 can recognize murine coronavirus mouse hepatitis virus (MHV) in brain macrophages, microglial cells, and oligodendrocyte cells.^17,18^ RIG-I also participates in the immune sensing of MHV in oligodendrocyte cells.^18^ Upon the sensing of cytosolic dsRNA by RIG-I/MDA-5, the adaptor protein MAVS (also known as VISA, Cardiff, or IPS-1) was recruited, which will activate TANK-binding kinase 1 (TBK1)/inhibitor of κB kinase epsilon (IKKε) and then induce the phosphorylation, and subsequent nuclear translocation of the transcription factor IFN regulatory factor 3 (IRF3), together with NF-κB that also activated by RIG-I/MDA-5 signaling, to initiate the transcription of type I and III IFNs and other proinflammatory cytokines, leading to antiviral immune responses.^14^ TLR3 also involves in the defense against SARS-CoV-1 infection.^19^ Activated TLR3 by dsRNA would activate IRF3 and NF-κB signaling via TIR-domain-containing adapter-inducing interferon-β (TRIF)-TBK1/IKKε cascades, resulting in the production of type I and III IFNs and other pro-inflammatory cytokines. ^14^ Although no evidence showed that the cytosolic DNA-sensing pathway of cGAS-stimulator of IFN genes (STING) signaling involved in recognition of coronaviruses, papain-like protease domain from SARS-CoV-1 could act as the antagonist of IFNs by targeting STING,^20,21^ suggesting that cGAS-STING pathway plays a vital role in defending against certain coronaviruses. STING was activated by the second messenger 2’-3’cGAMP produced by DNA-activated cGAS.^22^ Subsequently, STING recruits TBK1, which phosphorylates IRF3, leading to the trans-locations of IRF3 into the nucleus to turn on the expression of type I and III IFNs and inflammatory cytokines.^22^ The RIG-I/MDA-5-MAVS, TLR3-TRIF, and cGAS-STING signaling pathways converge on the points of TBK1/IKKε, which catalyzes IRF3 phosphorylation and the subsequent transcription of type I and III IFNs.^23^ Secreted type I and III IFNs bind to their receptors, respectively, and then activate Janus kinase/signal transducers and activators of transcription (JAK/STAT) signaling to drive the expression of IFN-stimulated genes (ISGs) which can initiate antiviral states by suppressing viral replication and spreading, activating immune cells, and causing cell deaths of infected cells.^15,24^

The type I and III IFN response is the essential action of host antiviral immunity in the clearance of virus infection. ^15,24^ To establish successful infection to host cells, viruses including coronaviruses have developed various strategies to antagonize the IFN response.^16^ The accessory proteins of SARS-CoV-1, such as open reading frame 3b (ORF3b), ORF6, and ORF9b have been proposed to inhibit the production of type I IFNs.^16^ In COVID-19 patients, it was observed that the induction of type I and III IFNs are suppressed.^9,10,25^ The replenishment of type I or III IFNs can significantly contribute to SARS-CoV-2 clearance and the relieve symptoms of COVID-19.^26-28^ Compared with type I IFNs, type III IFN shows certain advantages in the treatment of COVID-19 more for the induction of a more lasting antiviral stat and the less pro-inflammatory responses.^29^ Although it is observed that SARS-CoV-2 infection can impair the antiviral immunity mediated by type I and III IFNs in COVID-19 patients and cell models,^9,10,25^ for its recent emergency, how SARS-CoV-2 can block the induction of type I and III IFNs is still elusive. Therefore, dissecting the molecular mechanism of how SARS-CoV-2 evades human antiviral immunity elicited by type I and III IFN responses will contribute to understanding the pathogenesis of COVID19 and providing therapeutic strategies for treatments to counteract SARS-CoV-2 infections. In this study, we reported that an accessory protein of SARS-CoV-2, called ORF9b which is an alternative ORF within the *N* gene, can remarkable suppress type I and III IFN production induced by RIG-I/MDA-5-MAVS, TLR3-TRIF, and cGAS-STING signaling pathways by targeting multiple molecules of these innate antiviral signaling pathways.

## Materials and methods

### Reagents and antibodies

Protein A/G beads were purchased from Santa Cruz Biotechnology (USA), and the anti-Flag magnetic beads were purchased from Bimake. 2’3’-cGAMP was purchased from Invivogen. Rabbit anti-Myc-tag (71D10), rabbit anti-IRF3 (D83B9), rabbit anti-pIRF3 (4D46), rabbit anti-TBK1 (3031S), rabbit anti-pTBK1 (D52C2) were purchased from Cell Signaling Technology; Mouse anti-MAVS was purchased from Santa Cruz; Mouse anti-actin, mouse anti-V5-tag, and rabbit anti-calnexin were purchased from proteintech; Mouse anti-Flag M2 was purchased from Sigma Aldrich; Mouse anti-Myc-tag (9E10) was purchased from Origene; Rabbit anti-GM130 was purchased from Abcam; Rabbit anti-Tom20 antibody was purchased from Abclonal; Mouse anti-HA was purchased from MDL biotech.

### Constructs and plasmids

Plasmids expressing RIG-I, RIG-IN, MDA-5, MAVS, TBK1, IKKε, IRF3-5D, TRIF, and STING were cloned into mammalian expression vectors and the luciferase reporter plasmids including pGL3-IFN-β-Luc (IFN-β luciferase reporter) and pGL3-IFN-λ1-Luc (IFN-λ1 luciferase reporter) were constructed by inserting the promoter region into pGL3-Basic (Promega, USA) by standard molecular cloning methods as described in our previous publications.^30-32^ pISRE-Luc (the luciferase reporter of ISGs) was purchased from Clontech (USA). The SARS-CoV-2 ORF9b gene was synthesized according to the genome sequence of the SARS-CoV-2 Wuhan-Hu-1 strain (NC_045512.2) at General Biol (China). The coding region of the SARS-CoV-2 ORF9b gene was amplified using primers list in Supplemental Table 1 by PCR and cloned into the pCAG mammalian expression vector with a C-terminal Flag-tag.

### Cell culture

HEK-293, HEK-293T, HeLa, and Vero E6 cells were obtained from the American Type Culture Collection (ATCC), and cultured according to the culture method provide by ATCC. All these cells were cultured in Dulbecco’s modified Eagle’s medium (DMEM, Gibco, USA) with 10% heat-inactivated fetal bovine serum (FBS, Gibco, USA) at 37°C in a humidified incubator with 5% CO_2_.

### Transfection

The plasmids were transiently transfected into the cells using Lipofectamine 3000 (Invitrogen) or Polyethylenimine ‘Max’ (Polysciences, Inc., Germany) following the manufacturer’s instruction. Poly (I:C) and 2’-3’ cGAMP were delivered into cells using Lipofectamine 2000 (Thermo Fisher, USA) as described previously.

### RNA extraction and real-time quantitative PCR

Total RNA was extracted with TRIzol reagent (Invitrogen) and then was reverse-transcribed into first-strand cDNA with the HiScript III 1st Strand cDNA Synthesis Kit with gDNA wiper (Vazyme, China) following the manufacturer’s protocol. The SYBR Green-based RT-qPCR kit UltraSYBR Mixture (CWBIO, China) was used to perform real-time quantitative PCR (RT-qPCR) assays using primers of each gene (Supplemental Table 1) by a Roche LightCycler96 system according to the manufacturer’s instructions. The relative expression of the indicated genes was normalized to the mRNA level of GAPDH, one of the internal housekeeping genes in human cells. A comparative C_T_ method (ΔΔC_T_ method) was used to calculate the fold changes by normalizing to that of genes expressed in the control group as described previously.

### Dual-luciferase reporter assays

HEK-293T cells (approximately 0.5 × 10^5^/well) were seeded in 48-well plates. 12 hours before transfection. The luciferase reporter plasmids and the gene expression plasmids were co-transfected into HEK-293T cells as indicated in each figure legend. The pRL-TK Renilla luciferase reporter (Promega, USA) was also transfected to serve as an internal control. Thirty-six hours later, the cell lysate was used to assess the luciferase activities using the Dual-Luciferase Reporter Assay Kit (Vazyme, China) as described in our previous studies.^33-35^ The luciferase activity was measured in a Centro XS3 LB 960 microplate luminometer (Berthold Technologies, Germany). The relative luciferase activity was calculated by normalizing firefly luciferase activity to that of Renilla luciferase. The activity of firefly luciferase was normalized to that of Renilla luciferase to calculate the relative luciferase activity.

### Viral infection

VSV-enhanced green fluorescent protein (eGFP) and SeV were used to infect HeLa, HEK293, or HEK293T cells as described in our previous publications.^30-32^ Briefly, before infection, prewarmed serum-free DMEM medium at 37°C was used to wash the target cells, after which the virus was diluted to the desired multiplicity of infection (MOI) in serum-free DMEM and incubated with the target cells for 1-2 hours. At the end of the infection, the virus-medium complexes were discarded, and DMEM containing 10% FBS was added.

### Immunoblot analysis and immunoprecipitation

For Co-IP assay, HEK293T cells were first transfected with the indicated plasmids for 24 h and further lysed in lysis buffer [1.0% (v/v) NP-40, 50 mM Tris-HCl, pH 7.4, 50 mM EDTA, 0.15 M NaCl] complemented with a protease inhibitor cocktail (Sigma), and a phosphatase inhibitor cocktail (Sigma). Supernatants were transferred into new tubes after centrifugation for 10 min at 14,000g and further incubated with the indicated antibodies for 3 hours at 4 □ followed by the addition of protein A/G beads (Santa Cruz), or with Anti-Flag magnetic beads (Bimake), anti-Myc magnetic beads (Bimake). After incubation overnight at 4°C, beads were subject to washing four times with lysis buffer. After washing, the beads were boiled by boiling with 2×SDS loading buffer containing 100 mM Tris-HCl pH 6.8, 4% (w/v) SDS, 20% (v/v) glycerol, 0.2% (w/v) bromophenol blue, and 1% (v/v) 2-mercaptoethanol to collect the immunoprecipitates.

For immunoblot analysis, cell pellets were lysed with the M-PER Protein Extraction Reagent (Pierce) complemented with a protease inhibitor cocktail (Sigma) and a phosphatase inhibitor cocktail (Sigma). A bicinchoninic acid assay (Pierce) was used to measure the protein concentrations in the supernatant and protein samples were made equal with extraction reagent. The prepared total cell lysates or immunoprecipitates were electrophoretically separated by SDS-PAGE, transferred onto a polyvinylidene difluoride membrane (Millipore), blocked with 3% (w/v) bovine serum albumin (BSA), probed with indicated primary antibodies and corresponding secondary antibodies, and visualized by ECL Western blotting detection reagent (Pierce).

### Confocal microscopy

HeLa cells were seeded on 12-well slides 24h before transfection. Each well was transfected with the indicated plasmids (1µg each). Transfected or infected HeLa cells were subject to fix, permeabilization, and blocking as described in the previous paper. The fixation, permeabilization, and blocking buffer were all purchased from Beyotime Biotechnology. The cells were then incubated with indicated primary antibodies at 4°C overnight, rinsed, and incubated with corresponding secondary antibodies (Invitrogen). Nuclei were counterstained with DAPI (Abcam). Images were captured with a Zeiss LSM880 confocal microscope.

### Plaque assays

Vero-E6 cells were used to perform plaque assays to determine the titer of VSV-eGFP. Simply, Vero cells at approximately 100% confluency cultured in 24-well plates were infected with serial dilutions of VSV-eGFP. 0.5 hours’ later, the culture medium was discarded, and then DMEM containing 0.5% agar and 2% FBS overlaid. After 20 hours’ culture, the cells were fixed with a 1:1 methanol-ethanol mixture and then visualized with 0.05% crystal violet. The plaques on the monolayer were used to determine the titer of VSV-eGFP as described in our previous publications.^34^

### Statistical analysis

Statistical analysis was performed using two-tailed unpaired Student’s t-tests by GraphPad Prism 8.0 and Microsoft Excel. Unless specified, otherwise, one result, which is representative of three independent experiments, is chosen to be present as the mean ± SEM. The value of PD<D0.05 was considered to be statistically significant as indicated in each figure.

## Results

### SARS-CoV-2 ORF9b antagonizes type I and III IFNs

To explore the function of SARS-CoV-2 ORF-9b in viral infection, SeV was used to infect HEK293T cells expressing SARS-CoV-2 ORF9b. Using RT-qPCR analysis, we found that, compared with the control HEK293T cells that not express any viral protein, SARS-CoV-2 ORF9b expressing cells exhibit suppressed induction of IFN-β, IFN-λ1, and two ISGs called ISG56 and CXCL10 after SeV infection (Fig. 1). Similar results were observed when transfected a dsRNA mimic poly (I:C) into HEK293T cells to stimulate the antiviral immunity (Fig. 1). However, SARS-CoV-2 ORF-9c, another alternative ORF within the *N* gene, has no effect on neither SeV infection-nor poly (I:C) transfection-induced type I and III IFN production (Supplemental Fig. 1-2).

**Figure 1.**
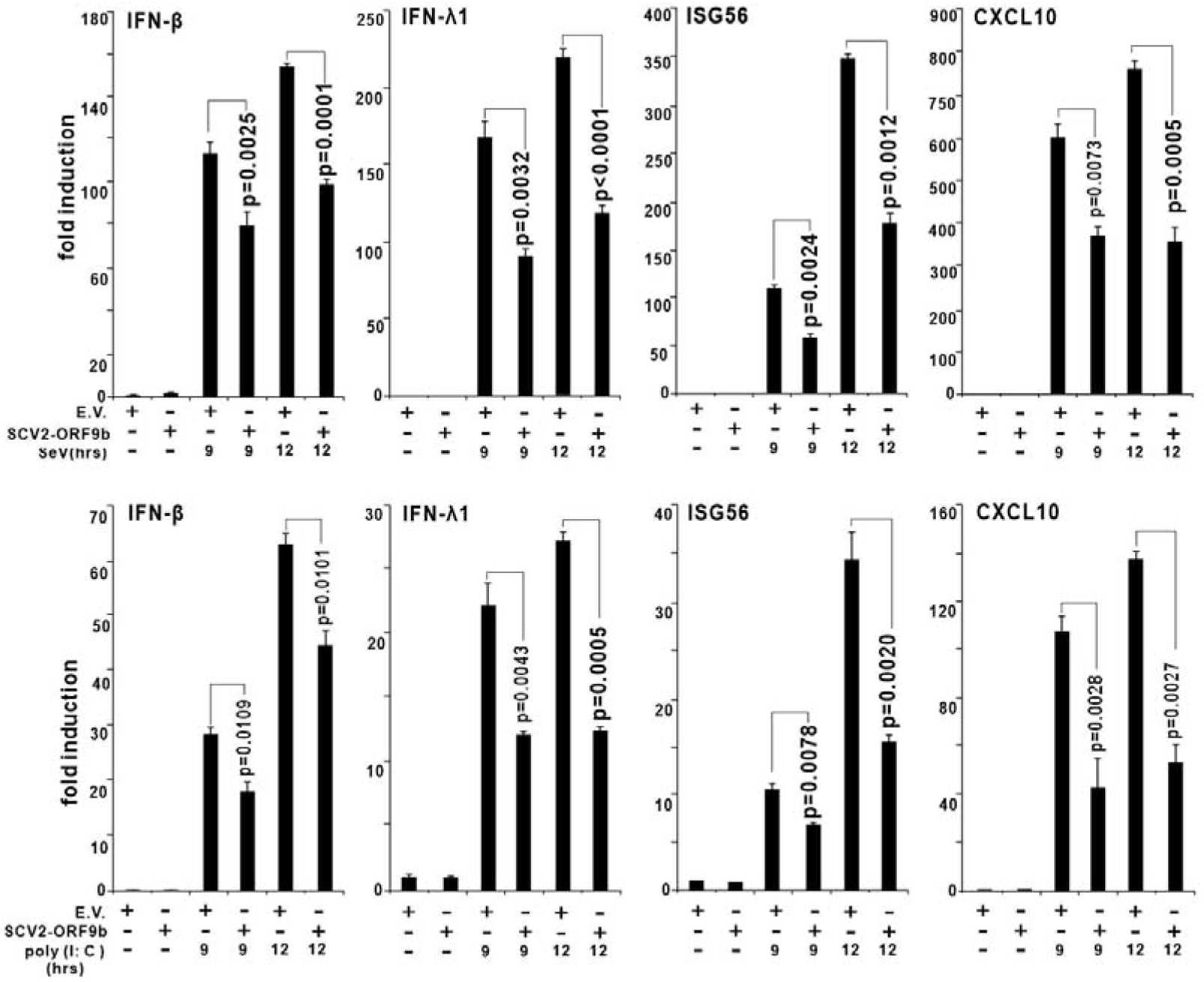
SARS-CoV-2 ORF9b suppresses the SeV- or poly (I:C)-induced IFN-β, IFN-λ1, ISG56, and CLXL10 production. Plasmids of pcDNA6B empty vector (500 ng) or SCV2-ORF9b (500 ng) were transfected into HEK-293T cells, 24 hours later, cells were infected with SeV infection or transfected with poly (I:C) transfection as indicated. At 9 and 12 hours after stimulation, the expression of IFN-β, IFN-λ1, ISG56, and CLXL10 in these cells was determined by RT-qPCR analysis. The results of one representative experiment was shown to represent three independent biological replicates. Error bars indicate SEM. empty vector: E.V; SARS-CoV-2 ORF9b: SCV2-ORF9b; hours: hrs.

Next, we try to map the layer where SARS-CoV-2 ORF-9b exerts its inhibitory effect on IFN production. We co-transfected SARS-CoV-2 ORF-9b with RIG-IN (an active form of RIG-I), MDA-5, MAVS, TBK1, and IRF3-5D into HEK293T cells. The luciferase reporter assays showed that SARS-CoV-2 ORF9b could obviously inhibit the activities of type I (IFN-β-Luc) and III (IFN-λ1-Luc) IFN and ISG (ISRE-Luc) luciferase reporters induced by RIG-IN, MDA-5, MAVS, TBK1, but not IRF3-5D, suggesting that SARS-CoV-2 ORF-9b inhibit RIG-I/MDA-5-MAVS signaling activated IFN production at the point upstream of IRF3 (Fig. 2). We also assessed the effect of SARS-CoV-2 ORF-9b on TLR3-TRIF and cGAS-STING signaling pathways by co-expressing SARS-CoV-2 ORF-9b with TRIF or STING. The results indicated that SARS-CoV-2 ORF-9b also suppressed TRIF- and STING-induced luciferase reporter activation of IFN-β, IFN-λ1, and ISGs (Fig. 2). Thus, it seems that SARS-CoV-2 ORF-9b exerts its IFN inhibitory function upstream of IRF3 but downstream of MAVS, TRIF, and STING.

**Figure 2.**
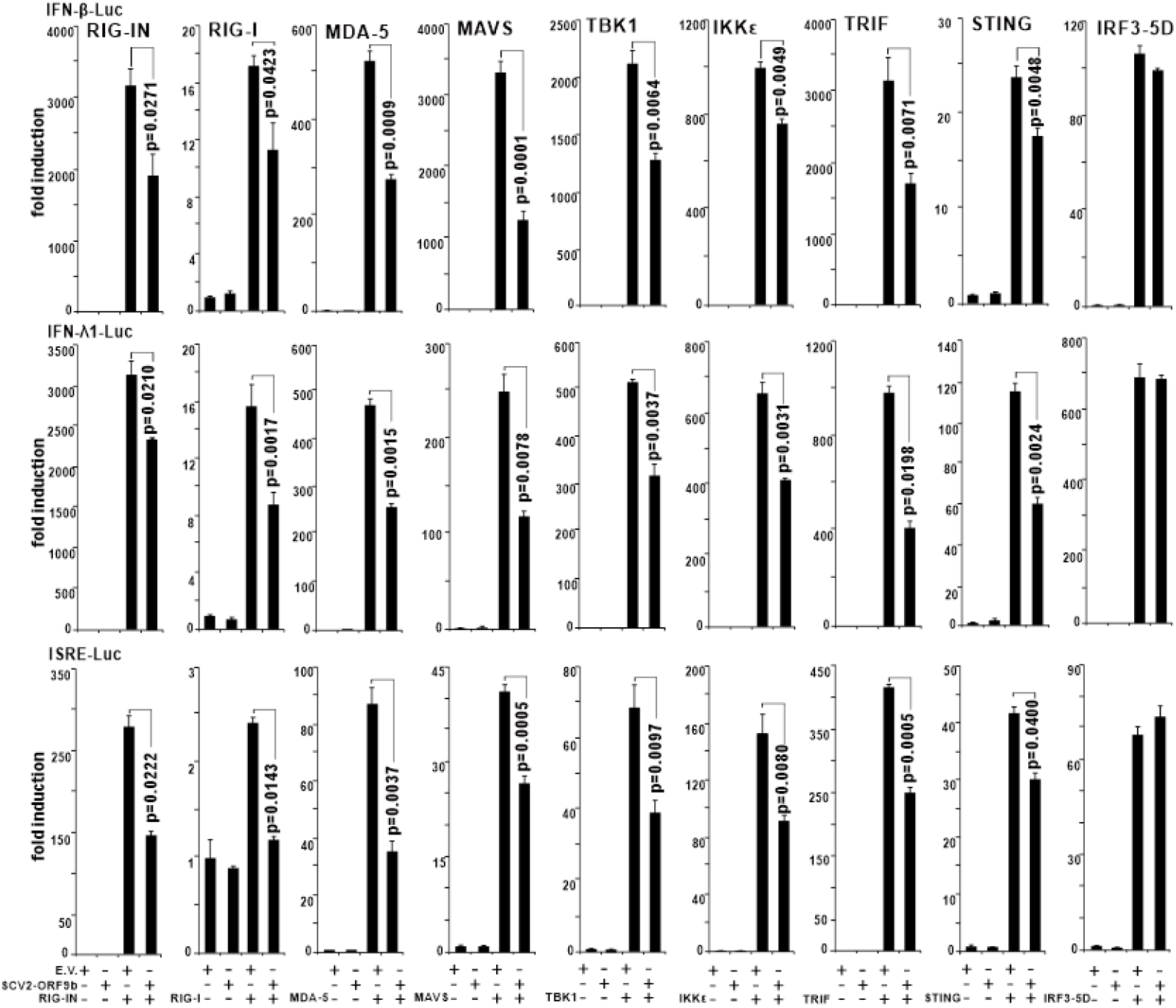
SARS-CoV-2 ORF9b inhibits the activation of luciferase reporters of type I and III IFNs and ISGs. Plasmids of RIG-IN (100 ng, an active form of RIG-I), MDA-5 (100 ng), TBK1 (100 ng), IKKε (100 ng), IRF3-5D (100 ng, an active form of IRF3), TRIF (100 ng, the adaptor of TLR3-TRIF pathway), or STING (100 ng, the adaptor of cGAS-STING pathway) were transfected alone or together with a plasmid expressing SARS-CoV-2 ORF9b into HEK-293T cells culture in 48-well plate as indicated. pRL-TK (5 ng) was also co-transfected to each well as an internal control. pcDNA6 empty vector was used to balance the total amount of plasmid DNA transfected into each well. Dual-luciferase assays were performed 36 h after transfection. Error bars indicate SEM. empty vector: E.V; SARS-CoV-2 ORF9b: SCV2-ORF9b.

### SARS-CoV-2 ORF9b targets multiple proteins of antiviral signaling pathways

To further localize the place where SARS-CoV-2 ORF9b performs its function, we first explore its subcellular localization using confocal microscopy. The plasmid expressing SARS-CoV-2 ORF9b was transfected into HeLa cells, 20 hours later, it was reacted with the primary antibody and then stained with fluorescent labeled secondary antibody as indicated (Fig. 3). The mitochondria, ER, and Golgi were visualized with corresponding markers. The results showed that SARS-CoV-2 ORF9b has a strong co-localization signal with mitochondria, but it is only weakly co-localized with ER and Golgi (Fig. 3). Next, we studied the co-localization of SARS-CoV-2 ORF9b with components of the innate antiviral signal pathways. Plasmids expressing RIG-I, MDA-5, MAVS, TBK1, TRIF, or STING was co-transfected with SARS-CoV-2 ORF9b plasmid into HeLa cells. The proteins were stained with florescent labeled secondary bodies after incubation with the primary antibodies as indicated. Confocal microscopy observation showed that SARS-CoV-2 ORF9b has a completely co-localization with MASV but partially localized with RIG-I, MDA-5, TBK1, TRIF, and STING (Fig. 3).

**Figure 3.**
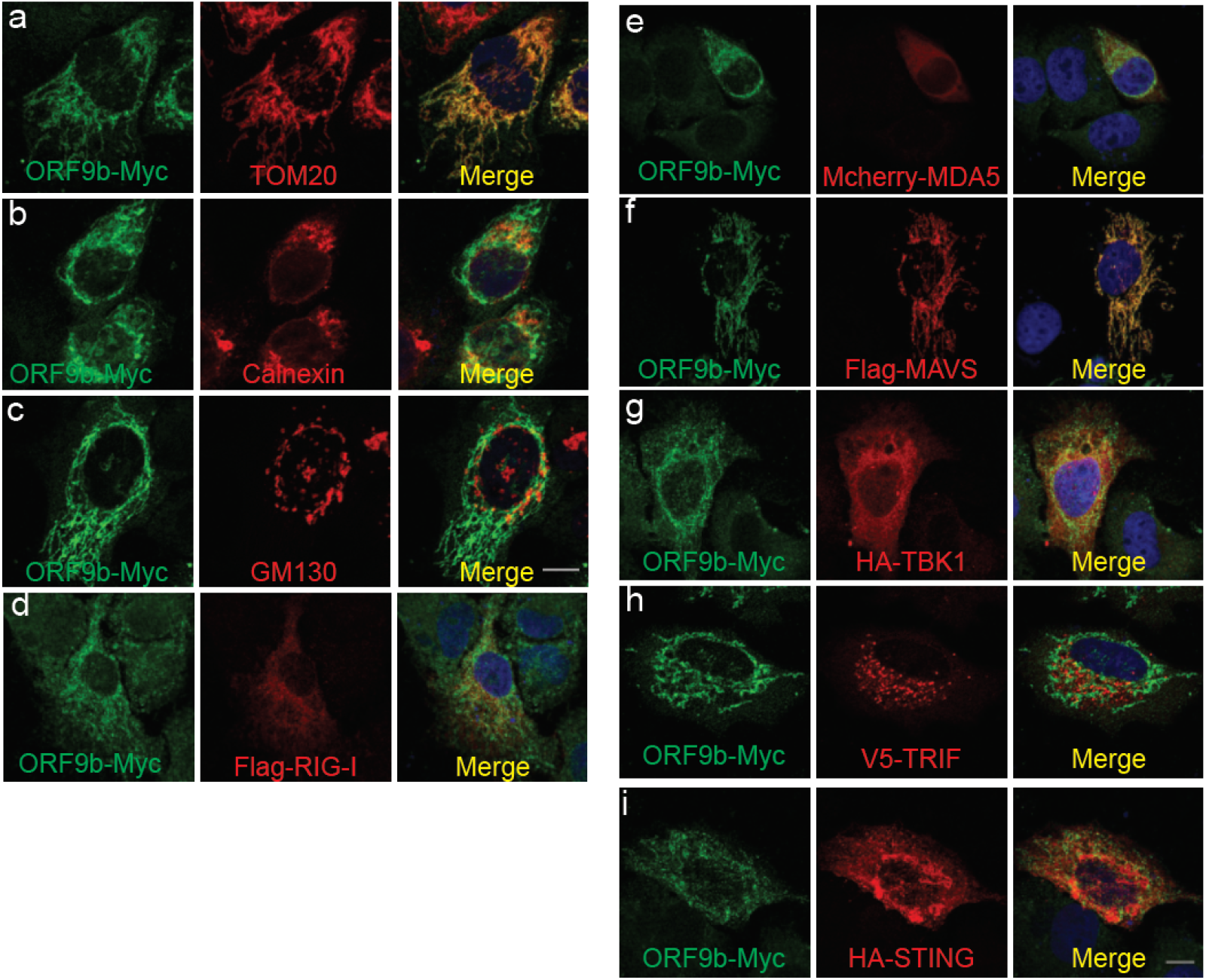
(a-c) Subcellular localization of SARS-CoV-2 ORF9b. HeLa cells seeded on 12 well coverslips were transfected with the indicated plasmids. After transfection for 20 h, HeLa cells were subject to immunofluorescence staining with mouse anti-Myc antibody and the rabbit antibodies against the corresponding organelle marker. Scale bar, 10 μm. (d) Relative localization of SARS-CoV-2 ORF9b protein with signaling molecules, including RIG-I, MDA5, MAVS, TBK1, TRIF, and STING. The seeding and transfection of HeLa cells were performed as same as in A). After transfection, ORF9b was stained with a rabbit anti-Myc antibody, and the signaling molecules were reacted with mouse antibodies against the indicated tags. Scale bar, 10 μm. TOM20, Mitochondria marker; Calnerxin, ER marker; GM130, Golgi marker. SARS-CoV-2 ORF9b protein, ORF9b.

Results from luciferase reporter assays suggest that the action step of SARS-CoV-2 ORF9b might be upstream of IRF3 (Fig. 2). We postulated that it might interact with molecules upstream IRF3 of the innate antiviral signaling pathways. To determine which protein is the target of SARS-CoV-2 ORF9b, we performed coimmunoprecipitation experiments. The plasmid expressing Myc-tagged SARS-CoV-2 ORF9b were co-transfected with plasmids of RIG-I, MDA-5, MAVS, TBK1, TRIF, STING, or IRF3 individually. Whole cell lysates were subjected to immunoprecipitation using antibodies as indicated (Fig. 4). Immunoprecipitation results indicate that SARS-CoV-2 ORF9b can associate with RIG-I, MDA-5, MAVS, TBK1, TRIF, and STING, but not IRF3, which consistent with the co-localization studies (Fig. 4). These data show that SARS-CoV-2 ORF9b may target multiple molecules of the RIG-I/MDA-5-MAVS, TLR3-TRIF, and cGAS-STING signaling pathways to suppress IFN production.

**Figure 4.**
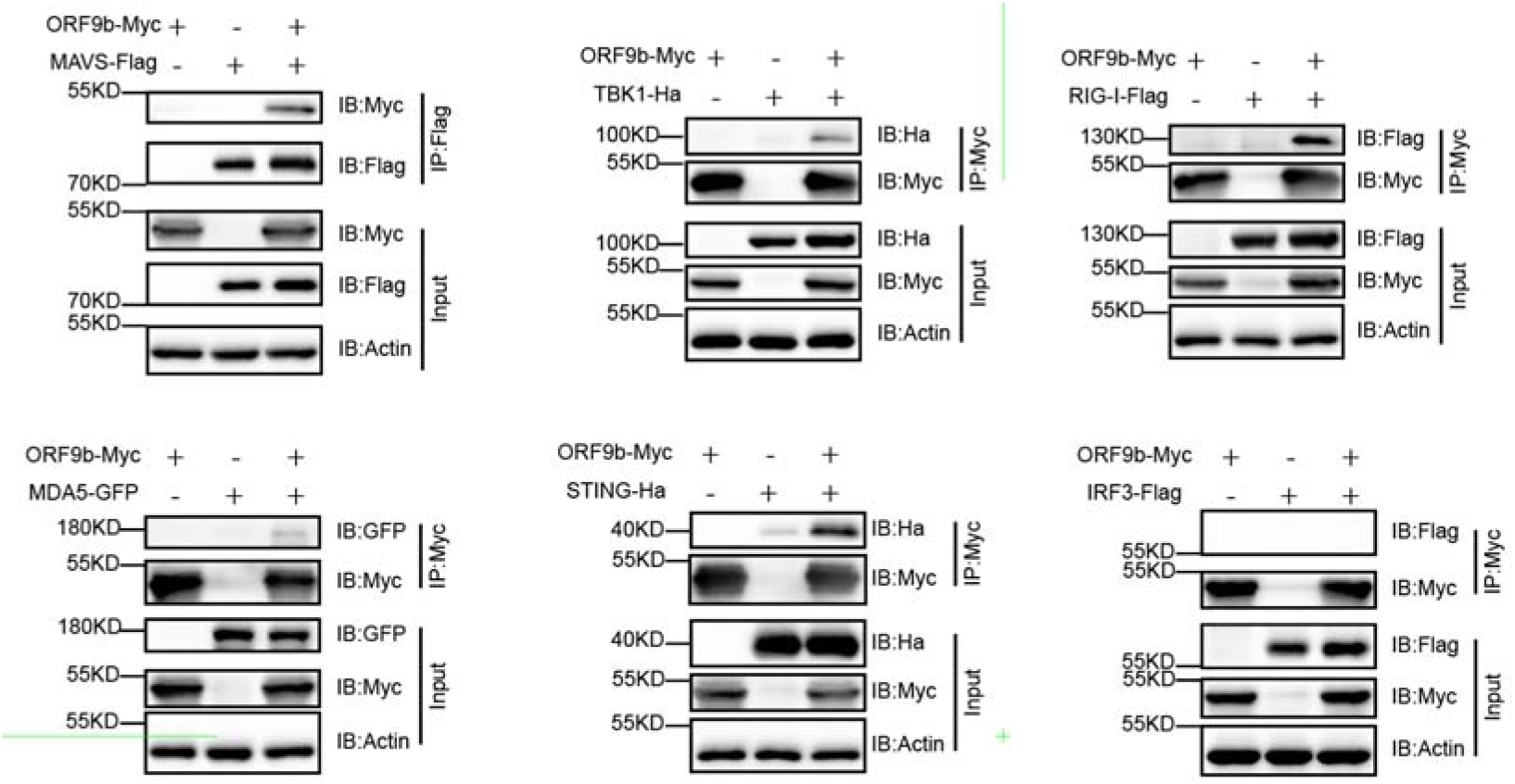
SARS-CoV-2 ORF9b interacts with RIG-I, MAVS, and TBK1 but not with IRF3. HEK293T cells were transfected with the indicated plasmids for 24h before co-immunoprecipitation by the indicated antibody-conjugated beads. The input and immunoprecipitates were reacted with the indicated antibodies.

### SARS-CoV-2 ORF9b decreases TBK1 phosphorylation

The signaling of RIG-I/MDA-5-MAVS, TLR3-TRIF, and cGAS-STING converge on the point of TBK1, SARS-CoV-2 ORF9b colocalizes and interacts with TBK1, thus the affection of TBK1 activity may be important to block signalling transduced from upstream molecules of RIG-I/MDA-5-MAVS, TLR3-TRIF, and cGAS-STING signaling pathways. Thus, we investigated the effect of SARS-CoV-2 ORF9b on TBK1 phosphorylation, an important biochemical process for IRF3 activation and subsequent IFN transcription. The phosphorylation of TBK1 was induced by overexpression of RIG-IN, MAVS, or TRIF, and by STING activated by its ligand 2’-3’cGAMP. When SARS-CoV-2 ORF9b plasmid was co-transfected, the phosphorylation of TBK1 induced by RIG-IN, MAVS, or STING but not by TRIF was dramatically reduced (Fig. 5). Thus, SARS-CoV-2 ORF9b can target TBK1 and prevented its phosphorylation induced by upstream signaling molecules of RIG-IN, MAVS, and STING but not TRIF. Further, we also observe that SARS-CoV-2 ORF9b can also impair TBK1 phosphorylation caused by SeV infection.

**Figure 5.**
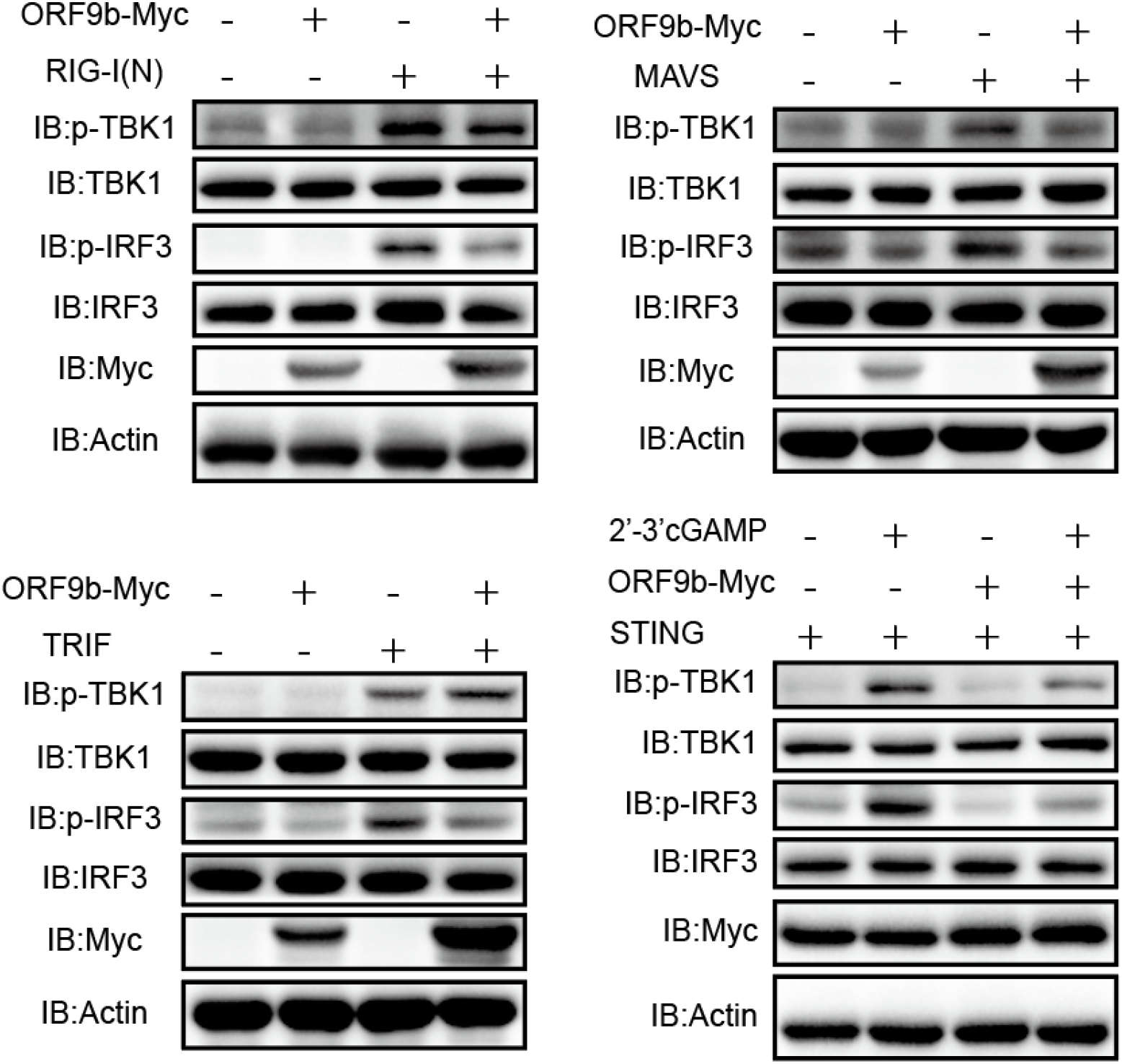
SARS-CoV-2 ORF9b suppresses TBK1 phosphorylation. (**a, b**) HEK293T cells were transfected with RIG-IN (a) or MAVS (b) in the presence or absence of ORF9b for 24h. (**c**) HEK293T cells were transfected with TRIF in the presence or absence of ORF9b for 24h. (**d**) HEK293T cells were transfected with STING in the presence or absence of ORF9b for 24h. The cells were further stimulated with transfection of 2’3’-cGAMP for 8h. The transfected or stimulated cells were lysed and subjected to immunoblot analysis with the indicated antibodies.

### SARS-CoV-2 ORF9b suppresses the phosphorylation and nuclear trans-localization of IRF3

TBK1 phosphorylation is a crucial step for IRF3 phosphorylation and nuclear translocation. Since SARS-CoV-2 ORF9b can block TBK1 phosphorylation induced by all three critical antiviral pathways of RIG-I/MDA-5-MAVS, TLR3-TRIF, and cGAS-STING signaling, it is of interest to determine whether SARS-CoV-2 ORF9b has any effect on the phosphorylation and nuclear trans-localization of IRF3. In HEK293T cells expressing RIG-IN, MAVS, TRIF, or STING, the phosphorylation of IRF3 was strongly induced, while, when SARS-CoV-2 ORF9b was co-expressed in these cells, IRF3 phosphorylation was dramatically impaired (Fig. 5). To determine whether SARS-CoV-2 ORF9b can still affect IRF3 phosphorylation in a viral infection, we used RNA virus, SeV, as a surrogate for SARS-CoV-2 to perform the virus infection studies. Control HeLa cells transfected with empty vector and HeLa cells expressing SARS-CoV-2 ORF9b were infected with SeV, 20 hours later, the cells were lysed for SDS-PAGE and immunoblotting analysis. The results indicated that SeV infection could induce the phosphorylation of both TBK1 and IRF3 (Fig. 6). When SARS-CoV-2 ORF9b was overexpressed in these cells, SeV-induced TBK1 and IRF3 phosphorylation was significantly suppressed. Thus, SARS-CoV-2 ORF9b can inhibit TBK1 and IRF3 phosphorylation stimulated by SeV infection.

**Figure 6.**
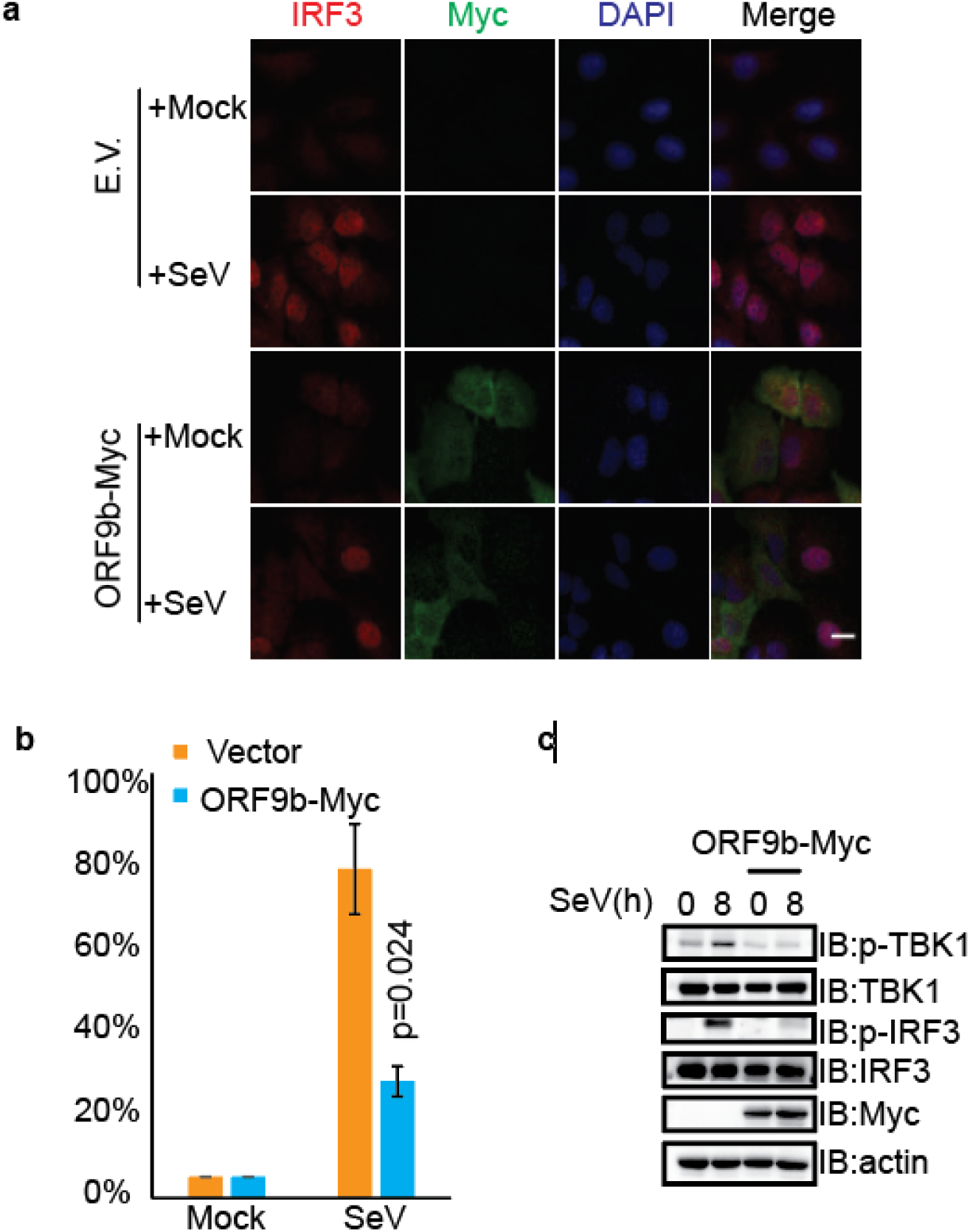
SARS-CoV-2 ORF9b suppressed IRF3 phosphorylation and nuclear translocation. (**a**) HeLa cells were seeded on 12 well coverslips (5*10^4 cells per well) one day before transfection. HeLa cells were further subjected to infection by SeV after transfection with the Myc vector plasmid or Myc-ORF9b plasmids for 20 h. Following infection for 8h, the slides were harvested and processed for immunofluorescence staining with mouse anti-Myc antibody and rabbit anti-IRF3 antibody. (**b**) Quantification of the percentage of IRF3 in the nucleus upon SeV infection. IRF3 localization from 50 cells within each group was counted and calculated before and after SeV infection. (**c**) SARS-CoV-2 ORF9b protein affects the phosphorylation of IRF3 upon SeV infection. HeLa cells seeded on 6 well plates (5*10^5 cells per well) were transfected with the Myc vector plasmid or Myc-ORF9b plasmid for 20 h before infection with SeV. At the indicated time points, cells were scraped and processed for immunoblotting with the indicated antibodies. SARS-CoV-2 ORF9b protein, ORF9b.

The transcription of IFNs was initiated by the phosphorylated IRF3 that translocated into the nucleus. The retention of IRF3 in the cytosol from its nuclear translocation will arrest its action on IFN induction. Since SARS-CoV-2 ORF9b inhibits IRF3 phosphorylation, next, we examined the effect of the SARS-CoV-2 ORF9b on SeV-induced IRF3 nuclear translocation. In resting cells, IRF3 was primarily distributed in the cytosol regardless of the expression of SARS-CoV-2 ORF9b or not (Fig. 6). When infected with SeV, IRF3 was translocated into nucleus of the control cells, however, IRF3 was restricted in cytosol in the cells expressing SARS-CoV-2 ORF9b (Fig. 6). This data supports that SARS-CoV-2 ORF9b can inhibit the nuclear translocation of IRF3 upon SeV infection.

### SARS-CoV-2 ORF9b impairs antiviral immunity

Although SARS-CoV-1 ORF9b suppresses IFN production, whether it can antagonize the antiviral immunity during a viral infection is still unknown. Thus, we activated the antiviral signaling pathways by transfecting TBK1, and explore whether SARS-CoV-2 ORF9b could affect the antiviral ability of cells expressing TBK1. HEK293T cells expressing TBK1 alone and HEK293T cells expressing both TBK1 and SARS-CoV-2 ORF9b were infected with VSV-eGFP, commonly used as a model virus to study the effect of IFN on viral replication. The infection of the virus was determined by examining the GFP positive cell under the fluorescent microscopy. The viral replication was determined by measuring the titer of the virus released into the culture medium. The results show that HEK293T cells expressing TBK1 and SARS-CoV-2 ORF9b have more GFP positive cells than that expressing TBK1 alone, suggesting that SARS-CoV-2 ORF9b might promote VSV-eGFP infection (Fig. 7). The viral titer from the culture medium of HEK293T cells expressing both TBK1 and SARS-CoV-2 ORF9b is higher than that from HEK293T cells expressing TBK1 alone; thus, SARS-CoV-2 ORF9b may facilitate VSV-eGFP replication (Fig. 7). These data indicate that overexpressing SARS-CoV-2 ORF9b can enhance virus infection and replication by blunting TBK1-induced antiviral immunity.

**Figure 7.**
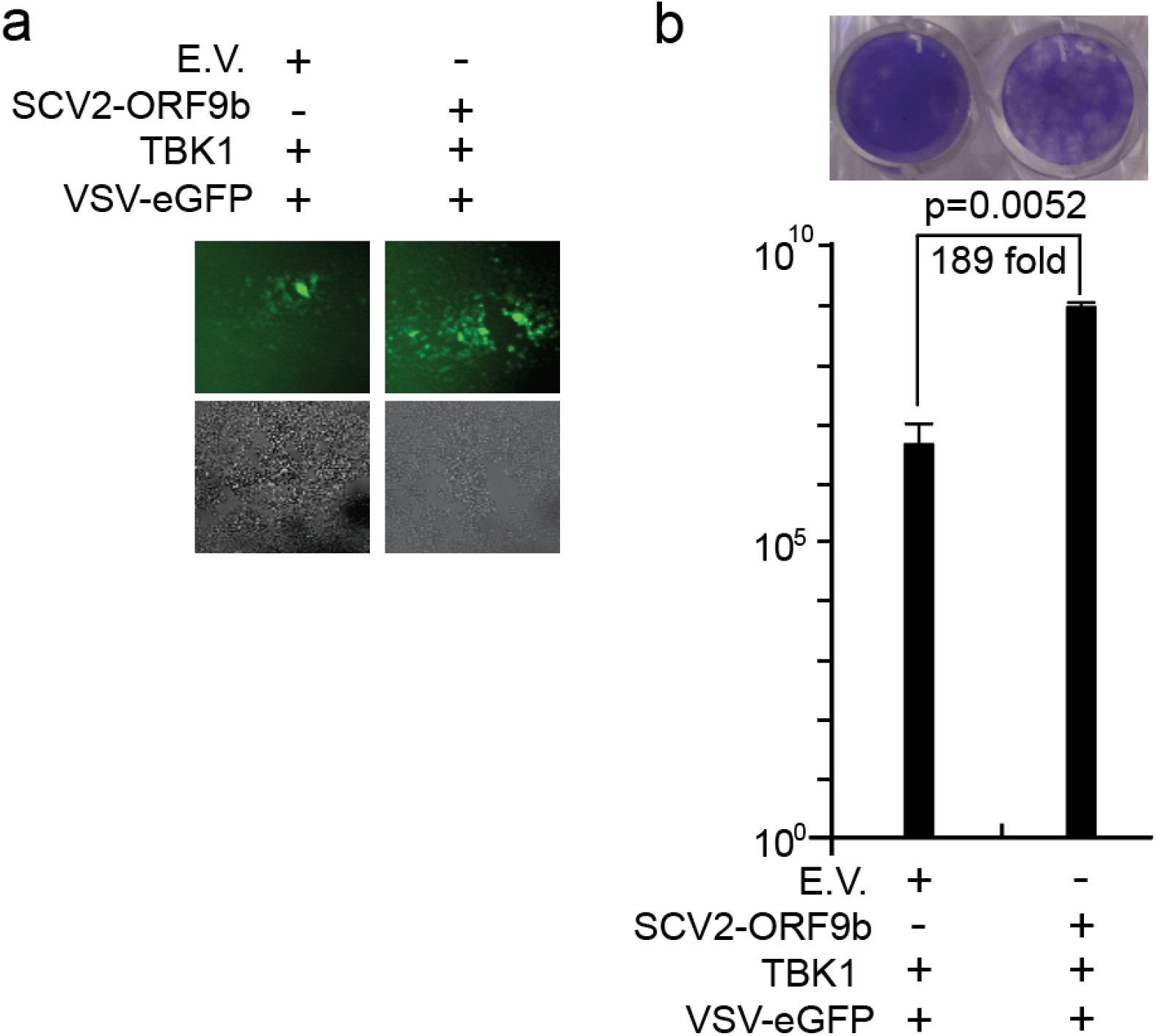
Overexpression of SARS-CoV-2 ORF9b impairs TBK1-dependent antiviral immunity. Plasmids were transfected into HEK-293 cells as indicated, 24 hours after transfection, the cells were infected with VSV-eGFP (MOI=0.001). Ten hours after infection, the GFP-positive cells were observed (**a**), and the culture supernatant (20 hours post-infection) was harvested for plaque assays to measure the titer of extracellular VSV-eGFP (**b**). Fluorescent imaging results are representative of two independent experiments. Scale bar, 50 μm. In panel b, the results of one representative experiment are shown, three independent biological replicates were analyzed, and the error bars indicate SEM. The statistical significance is shown as indicated. Empty vector, E.V.; SARS-CoV-2 ORF9b, SCV2-ORF9b.

## Discussion

The pandemic of COVID-19 is affecting the economy, transport, country’s relationships, and people’s life and health worldwide. A global effort is putting to find various strategies COVID-19 treatment. The immune system is essential for defense against virus infection; unfortunately, it is subverted by SARS-CoV-2 by suppressing the type I and III responses and enhancing the proinflammatory response, which will accelerate the viral replication and damage to host tissues and organs.^10^ Type I and III IFN responses play a critical role in human antiviral immunity against SARS-CoV-2 and restoration of type I and III IFNs in COVID-19 patients is effective as a therapeutic option in clinical trials, thus, how SARS-CoV-2 evade antiviral immunity is warranted.^15,27,28,36^ Here, we identify SARS-CoV-2 ORF9b can antagonize type I and III IFNs and impair host antiviral immunity. Overexpression of SARS-CoV-2 ORF9b inhibits the production of type I and III IFNs induced by SeV and poly (I:C). SARS-CoV-2 ORF9b inhibits the activation of RIG-I/MDA-5–MAVS, TLR3-TRIF, and cGAS-STING signaling pathways. Molecular mechanism studies show that SARS-CoV-2 ORF9b interacts with RIG-I, MDA-5, MAVS, TRIF, STING, and TBK1. SARS-CoV-2 ORF9b inhibits TBK1 phosphorylation induced by RIG-/MDA-5-MAVS and cGAS-STING signaling. Consequently, SARS-CoV-2 ORF9b inhibits the phosphorylation and nuclear translocation of IRF3 and type I and III IFN transcription. Furthermore, ectopic expression of the SARS-CoV-2 ORF9b protein facilitated the infection and replication of vesicular stomatitis virus. Thus, SARS-CoV-2 ORF9b can antagonize type I and III IFNs and contribute to the pathogenesis of COVID-19.

Among the coronaviruses, the homology genes of SARS-CoV-2 ORF9b only exist and was reported in SARS-CoV-1.^37^ SARS-CoV-1 ORF9b was characterized as an IFN antagonist by targeting mitochondria to enhances MAVS proteasomal degradation via its K48-linked ubiquitination. ^37^ SARS-CoV-1 and SARS-CoV-2 ORF9bs show 72.4% identity in amino acids. Surprisingly, one recently screening study shows that SARS-CoV-2 ORF9b does not affect IFN activation induced by RIG-I signaling.^38^ Thus, it is of great interest to further investigate whether SARS-CoV-2 ORF9b also involves in the suppression of IFNs. During the preparation of this manuscript, another recently showed that SARS-CoV-2 ORF9b suppresses type I interferon responses by targeting TOM70.^39^ Although the molecular mechanism of how Orf9b inhibits type I IFN responses through interacting with TOM70 is not investigated, it is proposed that ORF9b may compete with HSP90 for binding to TOM70 or may induce the production of lactic acid, which has been proven to inhibit IFN-I responses.^39^ Consistent with the above findings, we demonstrated that SARS-CoV-2 ORF9b inhibits the production of type I IFN induced by SeV infection, poly (I:C) stimulation, and the activation of RIG-I/MDA-5-MAVS, TLR3-TRIF, and cGAS-STING signaling pathways.

Lacking a biosafety level 3 laboratory, we have to use another RNA virus, SeV, to perform the viral infection studies. We found that overexpression of SARS-CoV-2 ORF9b can significantly reduce the production of IFN-β, IFN-λ1, ISG56, and CXCL10 stimulated by SeV infection (Fig. 1). We also found that SARS-CoV-2 ORF9b can inhibit SeV-induced phosphorylation and nuclear trans-localization of IRF3 (Fig. 6). Moreover, overexpression of SARS-CoV-2 ORF9b in HEK-293 cells can facilitate the replication of VSV-eGFP, which is sensitive to IFN signaling activation; thus SARS-CoV-2 ORF9b may enhance VSV-eGFP replication by suppressing IFN production (Fig. 7). Although SARS-CoV-1 ORF9b was reported to inhibit IFN production, its role in viral infection is currently unknown; thus, this is the first report that coronavirus ORF9b suppresses SeV-induced type I and III IFN production and promote VSV-eGFP replication.

Luciferase reporter assays showed that SARS-CoV-2 ORF9b could inhibit the promoter activities of IFN-β, IFN-λ1, and ISGs induced by multiple molecules of RIG-I/MDA-5-MAVS, TLR3-TRIF, and cGAS-STING signaling pathways (Fig. 2). We experimentally validated the interactions between SARS-CoV-2 ORF9b with numerous components of RIG-I/MDA-5 signaling pathways, such as RIG-I, MDA-5, and MAVS. Although we found that SARS-CoV-2 ORF9b associates with MAVS (Fig. 4) and inhibits MASV-induced IFN activation (Fig. 2), it cannot explain why SARS-CoV-2 also inhibit TRIF- and STING-induced IFN production. Thus, we hypothesis that SARS-COV-2 might (1) target TRIF and STING directly or target signaling molecules that parallel to or downstream of the converging point of RIG-I/MDA-5-MAVS, TLR3-TRIF, and cGAS-STING signaling pathways. Thus, firstly, we evaluate whether SARS-CoV-2 ORF9b directly targets TRIF and STING. Coimmunoprecipitation results indicated that SARS-CoV-2 ORF9b could be associated with TRIF and STING (Fig. 4), which also consists of the co-localization between SARS-CoV-2 ORF9b and TRIF or STING (Fig. 3). Although these results can explain why SARS-CoV-2 ORF9b can inhibit TRIF and STING-activated IFN signaling pathways, we are also curious about whether it can target TBK1, a molecular at the converging point of RIG-I/MDA-5-MAVS, TLR3-TRIF, and cGAS-STING signaling pathways. Coimmunoprecipitation results indicated that SARS-CoV-2 ORF9b could interact with TBK1 (Fig. 4); thus, demonstrated that SARS-CoV-2 ORF9b might perform its inhibitory effect on IFN production at the layer of TBK1. Combined with the results that SARS-CoV-2 ORF9b inhibits the induction of type I and III IFNs byTBK1 but not IRF3-5D, we propose that ORF8b fulfills this role at the layer or upstream of TBK1. Then we explore whether SARS-CoV-2 ORF9b can affect the phosphorylation of TBK1 and IRF3. Immunoblotting results showed that overexpression of SARS-CoV-2 ORF9b could inhibit TBK1 phosphorylation induced by RIG-IN, MAVS, STING, and SeV infection but not by TRIF (Fig. 5). These findings suggest that SARS-CoV-2 ORF9b inhibits RIG-I/MDA-5-MAVS signaling not only by interacting with RIG-I, MDA-5, and MAVS but also by perturbing TBK1 phosphorylation. Similarly, SARS-CoV-2 ORF9b suppresses cGAS-STING signaling by interacting with STING and inhibiting TBK1 phosphorylation. For the inhibition of TLR3-TRIF signaling, SARS-CoV-2 ORF9b can only target TRIF but does not have any effect on TRIF-induced TBK1 phosphorylation, suggesting that the inhibitory effect of SARS-CoV-2 ORF9b on TRIF-induced IFN signaling activation is achieved by interacting with TRIF directly but not by affecting TBK1 phosphorylation. In addition, the M protein of both SARS-COV-1 and 2 can inhibit IFN production by targeting RIG-I/MDA-5 signaling but have no effect on TBK1 phosphorylation;^34,40^ thus, the inhibition of TBK1 phosphorylation is not required for the antagonizing of IFNs by these viral proteins. Overexpression of SARS-CoV-2 ORF9b can inhibit IRF3 phosphorylation induced by RIG-IN, MAVS, STING, TRIF, and SeV infection, thus the inhibition of IRF3 phosphorylation may be indispensable for IFN antagonizing by these viral proteins. Therefore, SARS-CoV-2 ORF9b can target and interact with RIG-I, MDA-5, MAVS, TRIF, and STING, and impaired TBK1 phosphorylation aviated by the RIG-I/MDA-5-MAVS and cGAS-STING signaling pathways but not TLR3-TRIF signaling pathway. Overall, comparing with previous studies, we provide the following novel findings: besides MAVS, ORF9b may also associate with the dsRNA receptor RIG-I and MDA-5, TBK1, TRIF, and STING. Importantly, to our knowledge, we provided the first evidence that coronavirus ORF9b associate TBK1 and inhibit TBK1 phosphorylation induced by signaling from RIG-I/MDA-5, and cGAS-STING but not TLR3-TRIF signaling pathways.

The mitochondrial localization, where it degrades the MAVS signalosome, is essential for SARS-CoV-1 ORF9b to exert its IFN inhibitory function. We found that besides mitochondrial localization, SARS-CoV-2 also localizes on ER and Golgi (Fig. 3). ER is an important platform for TRIF and STING, while, Golgi is an important platform for and TBK1.^41-43^ Thus, these findings explain the co-localization and association of SARS-COV-2 ORF9b with TRIF, STING, and TBK1.The localization of SARS-CoV-2 ORF9b on ER and Golgi may provide the platform for its inhibition effects on TRIF, STING- and TBK1-induced IFN production. Thus, this study extended our understanding of the molecular mechanisms of coronavirus ORF6b-mediated IFN antagonizing.

SARS-CoV-2 is more sensitive to IFN treatment than other coronaviruses,^28^ multiple viral proteins become more critical to suppress IFN production at different steps to ensure the production and function of IFNs are minimized during SARS-CoV-2 infection. SARS-COV-2 ORF9b targets multiple proteins of the distinct immune signaling pathways, which may suppress IFN signaling at different steps. Similarly, SARS-CoV-2 ORF6 and MERS-CoV ORF4b are capable of perturbing multiple innate antiviral signaling pathways by target various components of these pathways.^38,44,45^ Although cGAS-STING is cytosolic dsDNA sensing pathway, coronaviruses, a family of RNA viruses, also encode viral proteins such as papain-like protease to impair STING function; thus, this pathway is essential in defending against coronavirus infection.^20,21^ And the inhibition of cGAS-STING pathway by SARS-CoV-2 ORF9b may suggest that this pathway may play a role in SARS-CoV-2 clearance, thus, drugs or chemicals such as 2’-3’cGAMP that activates this pathway may be considered to be used in COVID-19 treatment.

Distinct methods have shown that SARS-CoV-2 ORF9b inhibits type I and III IFN production by targeting multiple proteins of antiviral signaling pathways, we should be conscious that the transfection system might be different from the real viral infection. Therefore, further studies should be conducted in the context of real SARS-CoV-2 infection experiments. The accessory proteins of coronaviruses have been proposed to be not essential for viral replication. Ther are not directly involved in viral assembly,^7^ thus, theoretically, the *ORF9b* null SARS-CoV-2 might available if it cannot affect the translation and expression of N protein which is a structural protein for virion assembly. However, whether this *ORF9b* null SARS-CoV-2 virus is accessibility still need experimental validation. When this mutant virus is available, the real viral infection studies should contribute to our understanding of the role of ORF9b in IFN antagonizing.

Although the administration of exogenous IFNs is show to be valid for SARS-CoV2 clearance on both SARS-CoV-2 patients and cell models,^26,29,36,46^ full evaluation of this treatment requires extensive studies on the relative importance of all IFN-antagonizing viral proteins encoded by SAR-CoV-2. Thus, our findings that the SARS-COV-2 ORF9b suppresses type I and III IFN production contribute to our understanding of the pathogenesis of COVID-19, and the identification of multiple protein targets may provide more precise treatment of COVID-19.

## Acknowledgements

This work was supported by grants from the COVID-19 emergency tackling research project of Shandong University (Grant No. 2020XGB03 to P.-H.W), grants from the Natural Science Foundation of Jiangsu Province (SBK2020042706 to P.-H.W), grants from the Natural Science Foundation of China (81930039, 31730026, 81525012) awarded to C.G, and the Fundamental Research Funds of Shandong University (21510078614099), the Fundamental Research Funds of Cheeloo College of Medicine (21510089393109), China Postdoctoral Science Foundation (2018M642662), and the Natural Science Foundation of China (81901604) awarded to Y.Z. We thank Translational Medicine Core Facility of Shandong University for consultation and instrument availability that supported this work.

## Contributions

C.G. and P.-H.W. conceptualized the study. L.H., M.-W.Z., Y.Z., J.Z., and M.L., N. performed the experiments. P.-H.W. wrote the first draft of manuscript. All of the authors contributed to revising the manuscript and approved the final version for publication.

## Supplemental materials

**Supplemental Figure 1.**
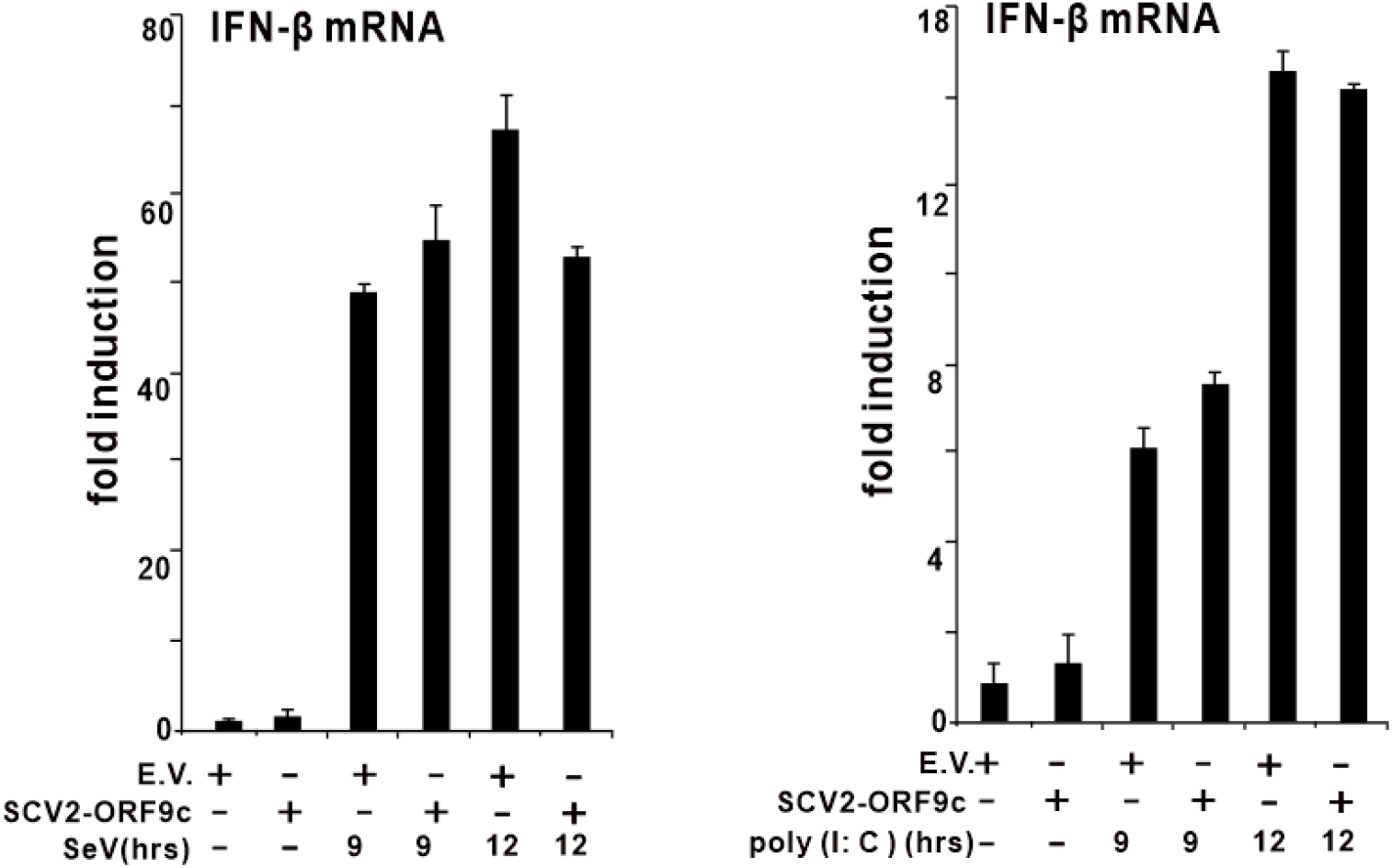
SARS-CoV-2 ORF9c has no effect on IFN-β induction by the stimulation of SeV infection and poly (I:C transfection. The plasmids of pcDNA6B empty vector (500 ng) or SARS-CoV-2-ORF9c (500 ng) were transfected into HEK293T cells cultured in 24 well plates. The cells were stimulated, and the expression of IFN-β, IFN-λ1, ISG56, and CLXL10 were determined similar to those described in figure 1. empty vector: E.V; SARS-CoV-2 ORF9c: SCV2-ORF9c; hours: hrs.

**Supplemental Figure 2.**
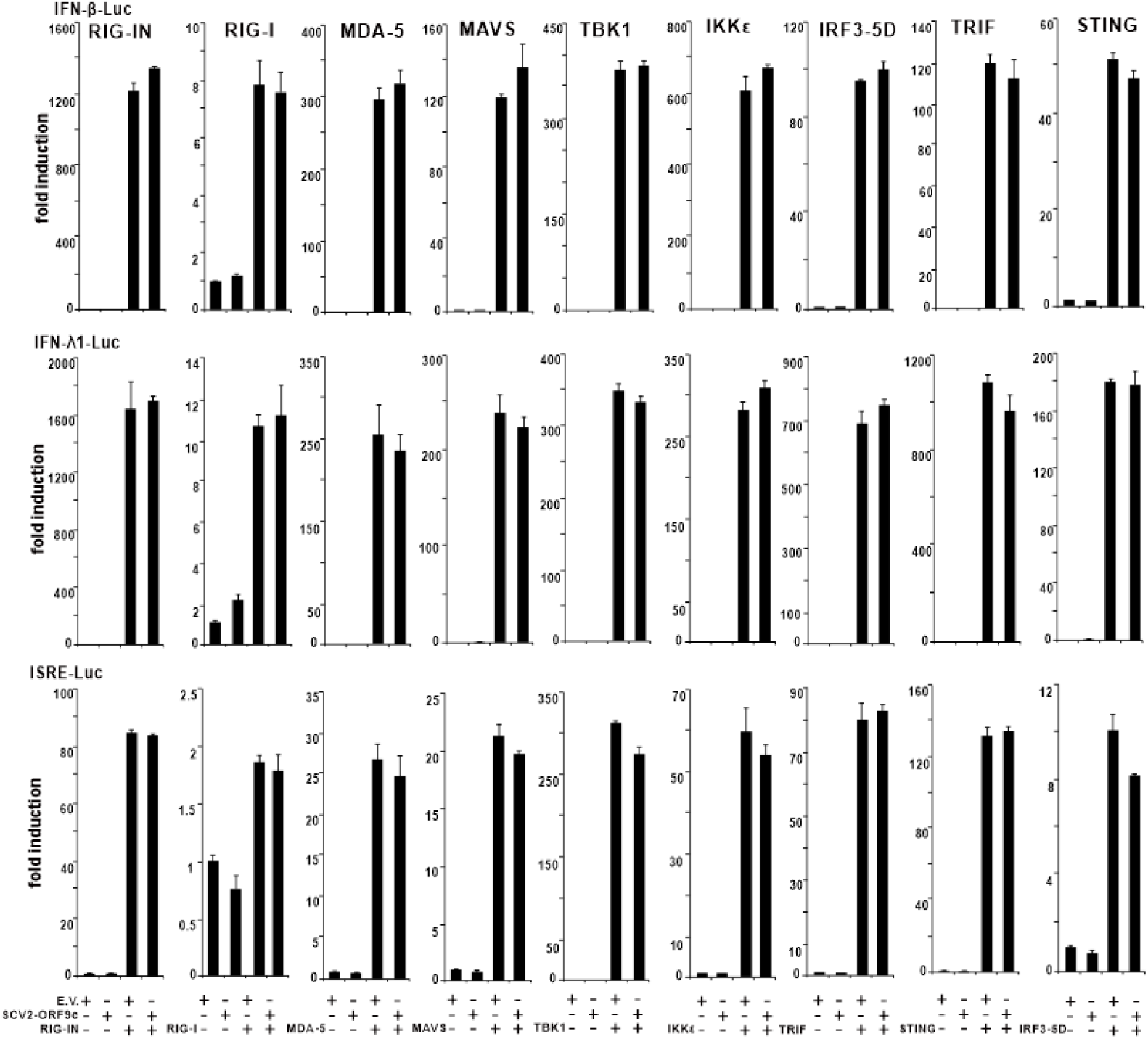
SARS-CoV-2 ORF9c does not have any effect on the activation of luciferase reporters of type I and III IFNs and ISGs. The combination of plasmids as indicated were transfected into HEK-293T cells as described in figure 2. 36 hours after transfection, dual-luciferase assays were performed. empty vector: E.V; SARS-CoV-2 ORF9c: SCV2-ORF9c.

